# Genomic signatures of local adaptation in marram grass highlight the importance of provenance in dune restoration

**DOI:** 10.64898/2026.11.06.764831

**Authors:** Emily R Baker, Emma V Curran, Carlijn Lammers, Emily J Bailes, Rachel Patel, Lucy S Knowles, Samuel GS Hibdige, Sahr Mian, Maria S Vorontsova, Ilia J Leitch, Josien Grashof, Emma Wolff, Laurence Jones, Nicola J Nadeau, Valérie C Reijers, Luke T Dunning

**Author notes:** These authors contributed equally to the work (Emily R Baker, Emma V Curran). Corresponding author: Luke T. Dunning; Ecology and Evolutionary Biology, School of Biosciences, University of Sheffield, Western Bank, Sheffield S10 2TN, United Kingdom; +44 equal(0) 1142220027.

## Abstract

Coastal sand dunes provide critical natural flood defences and key habitats for a range of threatened species. These systems are formed when shifting sands are stabilised by specialised coastal grasses. In Europe, marram grass (*Calamagrostis arenaria*) is the main dune building species. However, despite its ecological importance and its widespread use in dune restoration, little is known about the genomic innovations that facilitated the colonisation of the coast, the extent of local adaptation in current populations, or the genetic consequences of restoration practices. Using a chromosome-scale reference genome we generate here, we show that *C. arenaria* has significant expansions in several gene families relevant to coastal adaptation, including those associated with salinity, drought, and nutrient limitation. Using ddRAD-Seq data for over 200 individuals from 10 European populations, we identified strong genetic differentiation between northern and southern populations which corresponds to recognised subspecies. Within the northern european populations, the genetic structure is largely driven by isolation-by-distance. Although, we detected evidence of non-local planting in a restored Swedish dune system, where introduced plants were genetically distinct from the native population and both groups exhibited reduced genetic diversity. Genome-wide scans for local adaptation within the northern European marram grass individuals identified outlying genomic regions that were enriched for genes with functions associated with nutrient deficiency, drought, pathogen defence, and other environmental stressors characteristic of coastal dunes, potentially indicating local adaptation. Outlying regions also include genes associated with lignin biosynthesis and culm mechanical strength, potentially reflecting selection on the structural properties required to withstand different burial and wind exposure regimes. Overall, these findings show widespread local adaptation in *C. arenaria*, and support the use of local provenance in dune restoration and conservation.

## Introduction

Understanding the processes that shape genetic variation across populations and landscapes is not only a central goal of evolutionary biology, but it is crucial in guiding efforts to manage natural populations (Slatkin, 1987; Manel *et al*., 2003; Hohenlohe, *et al*., 2021). Characterising how genetic variation is structured within and between populations allows us to assess the relative importance of evolutionary processes such as gene flow, selection, and genetic drift on populations, which can directly inform management practices (Hohenlohe, *et al*., 2021). The extent of connectedness of populations via gene flow can impact how vulnerable populations are to size fluctuations, inbreeding depression and local extinctions (Hoffmann *et al*., 2021). Understanding the extent of local adaptation across the range of a species of interest is important when sourcing individuals to bolster populations of concern. For example, in revegetation schemes, introducing plants of local provenance can help maintain locally adapted alleles and prevent outbreeding depression, whereas introducing non-local alleles may help mitigate the effects of climate change (Breed *et al*., 2013). In some cases ecological restoration has been shown to decrease genetic diversity of populations (Wei *et al*., 2023). It is therefore important that we examine the genetic diversity of managed and unmanaged populations to assess the impact of management practices.

Coastal dunes are among the most intensively managed natural habitats in Europe. Shifting dunes can be stabilised by the widespread replanting and translocation of dune-building species, with the aim to mitigate erosion and flooding under anthropogenic climate change (Fernández-Montblanc, Duo and Ciavola, 2020; Gao et al., 2020; Lithgow et al., 2013). Dunes are highly dynamic and spatially heterogeneous environments (Hesp and Martínez, 2007; Bar, Katz and Dorman, 2023), exposed to strong abiotic stressors: salinity, drought, nutrient limitation, wind exposure, sand burial and frequent physical disturbance (Gilbert, Pammenter and Ripley, 2008; Hesp, 1991). Selective pressures can vary markedly over short temporal and spatial scales, generating steep environmental gradients that may promote local adaptation (Hesp, 2002; Du and Hesp, 2020).

Marram grass (*Calamagrostis arenaria* (L.) Roth, formerly *Ammophila arenaria*) is the dominant dune-building species in Europe and has been the subject of extensive replanting for more than a century as part of dune management and restoration programs worldwide (Huiskes, 1997; Perrino et al., 2013; Bonte et al., 2021; Rozé and Lemauviel, 2004; De Lillis, Costano and Tinelli, 2004). Through extensive clonal growth via rhizomes and rapid vertical elongation in response to sand burial, marram grass traps sediment and facilitates foredune development providing critical protection against coastal erosion and storm surges (Maun and Lapierre, 1984; Borsje et al., 2011; Temmerman et al., 2013). The species life history combines clonal propagation with sexual reproduction via wind pollination, alongside dispersal by both seeds and vegetative fragments, including the potential for long-distance transport by marine currents (Van Der Putten, 1990; Konlechner and Hilton, 2009; Lammers et al., 2023). These traits raise questions about levels of genetic diversity and how genetic variation is structured, given the potential for gene flow (both natural and human-mediated) and local adaptation.

Despite its ecological and societal importance, the population genetic structure of marram grass across Europe remains poorly resolved at the genomic level. Two subspecies are recognised: the northern *C. arenaria* subsp. *arenaria* and the southern *C. arenaria* subsp.. *australis* (Mabille) Asch. & Graebn. which are morphologically distinct and broadly separated geographically, with their ranges meeting in the French Basque Country (Tutin, 1980; Bogdanovic, Segota and Alegro, 2018). Previous studies using limited genetic markers have demonstrated differentiation between northern and southern populations, consistent with subspecific divergence, but have lacked the resolution to characterise fine-scale population structure, admixture and connectivity within regions (Rodríguez-Escheverría, Freitas and Van der Putten, 2008; Hol, van der Wurff, Skøt and Cook, 2008). Lack of genomic resources has also limited our understanding of the adaptations which help *C. arenaria* succeed in the harsh dune environment, and the extent to which the heterogeneous nature of the dune habitat combined with the broad latitudinal gradient encompassed by the species’ range, has driven local adaptation across populations.

Here, we combine population and comparative genomics approaches to examine the dynamics of population genetic structure in *C. arenaria,* alongside investigating both species-level, and population-specific (i.e. local) adaptation to the dune environment on a genome-wide level for the first time. We generate a chromosome-scale reference genome to (i) detect gene family expansions and contractions associated with coastal adaptation. We then use restriction site–associated DNA sequencing (RAD-seq) to (ii) characterise the population structure of *C. arenaria* across its Western European range, and assess the contributions of gene flow, subspecific isolation, and dune management activity in shaping it. Finally, we use a genome scanning approach to (iii) test for evidence of local adaptation across a latitudinal cline in *C. arenaria* spp. *arenaria.* This study both provides fresh insight into coastal adaptation, the evolutionary dynamics of an ecologically and societally important species, as well as providing practical implications for the provenancing of plants for dune restoration efforts.

## Materials and methods

### Generating a reference genome

A *de novo* reference genome was generated for a *C. arenaria* accession (UK-SFB-005) collected in 2021 from Saltfleetby-Theddlethorpe Dunes National Nature Reserve, United Kingdom (53°22’15.3“N, 0°14’26.9”E). The 1C genome size was estimated by flow cytometry using the one-step protocol (Doležel et al., 2007) with minor modifications (see Clark et al., 2016). *Petroselinum crispum* ‘Champion Moss Curled’ (2C = 4.50 pg; Obermayer et al., 2002) was used as the calibration standard and nuclei were isolated in the General Purpose Buffer (GPB; Loureiro et al. 2007), supplemented with 3% (w/v) PVP and 0.08% (v/v) beta-mercaptoethanol.

DNA was extracted after the plant was dark treated for 24 hrs. In total, 3 g of young leaf material was flash frozen with liquid nitrogen, ground using a pestle and mortar, and stored at -80 °C prior to DNA extraction. Total DNA was extracted using the NucleoBond HMW DNA kit (Macherey-Nagel), following the manufacturer’s protocol. DNA extractions were purified using AMPure PB Beads (Beckman Coulter Life Sciences), and purity, concentration and quality was assessed with NanoDrop 8000 (Thermo Fisher Scientific), Qubit 3 (Thermo Fisher Scientific) and Femto Pulse (Agilent), respectively. PacBio DNA libraries were prepared and run on eight Sequel II SMRT Cells in CCS mode at the NERC Environmental Omics Facility (NEOF) Liverpool (University of Liverpool, UK).

For the PacBio HiFi reads (QV ≥20), we estimated the coverage and ploidy of the UK-SFB-005 accession using the k-mer spectrum generated with GenomeScope v2.0 and Smudgeplot v.0.5.3 (Ranallo-Benavidez et al., 2020) with default parameters. The PacBio HiFi reads were *de novo* assembled using hifiasm v.0.16.1 (Cheng et al., 2021), with default parameters except for the number of haplotypes being set to four. Protein coding genes were annotated using Helixer v0.3.0 (Holst et al., 2025) with the default parameters and the recommended land plant model (land_plant_v0.3_a_0080.h5). The completeness of the assembly and annotation were assessed using BUSCO v3 (Waterhouse et al. 2018) with the embryophyta_obd10 database. The chloroplast genome was independently *de novo* assembled from the HiFi data using the oatK toolkit v.1.0 (with parameters -k 1001 -c 350; Zhou et al. 2025).

### Gene family expansion and contractions

To identify gene family expansions and contractions we used the OrthoVenn3 web server (Sun et al. 2023), with the predicted proteomes of *C. arenaria* and four other tetraploid grasses to reduce the effect of differences in ploidal level on the analysis (*Oryza coarctata* [BioProject PRJNA598408; Bansal et al., 2021], *Lolium Perenne* [BioProject PRJNA222646; Nagy et al., 2022], *Triticum turgidum* [BioProject PRJEB22687; Maccaferri et al., 2019], and *Eleusine coracana* [BioProject PRJNA838475; Devos et al., 2023]). For the OrthoVenn3 analysis we used the OrthoMCL (Li, Stoeckert and Roos, 2003) algorithm to identify orthologous proteins among the five species, CAFE5 (Mendes et al., 2020) to estimate which gene families are expanding or contracting, and a Gene Ontology (GO) enrichment analysis to determine whether these changes were significantly associated with specific functions or biological processes.

### Population level sampling and sequencing

Silica preserved leaf material was collected from multiple individuals from 10 sites across 6 European countries (Belgium, Denmark, The Netherlands, Portugal, Sweden, and UK) in 2022 (Table S1). Where possible, samples were collected from different stages of the dune system (i.e. the beach, yellow dunes [foredunes], and grey dunes further inland). DNA extractions were performed using the 96 Well SYNERGY Plant DNA Extraction Kit (OPS Diagnostics) following the manufacturer’s protocol. DNA extraction quantity and purity was assessed by NanoDrop and Qubit, and sent for restriction-site associated DNA (RAD) library preparation and sequencing (Peterson et al. 2012) at IGATech (Udine, Italy). Briefly, the DNA of each sample was double digested using 2.4 U of *PstI* and *EcoRI* endonucleases (New England BioLabs), barcodes were ligated to the resulting fragments, and the products were pooled prior to sequencing with NovaSeq 6000 (Illumina, San Diego, CA).

### Population structure

The RAD-seq data was demultiplexed, remaining adaptors trimmed, and low-quality reads (where >40% of bases have a phred quality score <20) removed using fastp v.0.24.0 (Chen et al. 2018) with default parameters. To remove chloroplast RAD-seq data, we mapped the cleaned reads to the UK-SFB-005 chloroplast assembly using RADOrgMiner v.0.95 (Laczkó et al. 2022) with default parameters. The unmapped nuclear RAD-seq reads were then used for downstream analysis, and were aligned to the genome assembly using BWA-mem v.2.2 (Vasimuddin et al. 2019) with default parameters. We removed samples with insufficient quantity (< 1 million reads) or quality (<80% of reads mapped) from the dataset.

Genotype likelihoods were estimated using ANGSD v.0.940 (Korneliussen et al. 2014) for sites present in ≥50% individuals, with ≥ 5 reads per individual, and mapping quality ≥20. Population structure was examined with principal component analysis (PCA) to summarise genetic variation across the samples, using PCAngsd v1.36.4 (Meisner and Albrechtsen 2018) and eigenvector decomposition in R v4.4.3. Genetic clusters and admixture proportions were estimated from the genotype likelihoods with NGSadmix v3.2 (Skotte et al. 2013), fitting between 2 and 10 genetic clusters (*K*), with 10 replicates for each *K*. The best fitting value for *K* was selected using the Δ*K* method (Evanno et al. 2005; Cullingham et al. 2020) implemented using CLUMPAK (Kopelman *et al*., 2015).

### Isolation-by-distance

Isolation-by-distance between the 10 populations was estimated by comparing the genetic distance and geographic distance between populations. Single nucleotide polymorphisms (SNPs) were called from the BWA-mem alignments using bcftools v.1.15.1 (Li, 2011) with default parameters. Sites with <5 reads per individual were marked as missing, and sites with >20% missing data across individuals and a quality score <20 were filtered out, and finally we only retained biallelic SNPs. Pairwise genetic distances were calculated between all populations using the Hudson’s F_ST_ estimator, implemented in plink v1.90 (Chang *et al*., 2015). The population from Ystad in Sweden was split into two for this analysis as two strongly distinct genetic clusters within this population were identified in the PCA (see results). The straight-line geographic distance between populations was calculated using the haversine method with the R package geosphere (Hijmans *et al*., 2025). The correlation between the genetic and geographic distance matrices was tested using Mantel tests with 10,000 permutations. A phylogenetic tree of all samples was also inferred using the SNP calls. A maximum likelihood tree with the software RAxML v.8.2.12 (Stamatakis, 2014) with a GTRCAT model and 100 bootstrap replicates.

### Genetic diversity

Genetic diversity (**π**) and Tajima’s D were measured within each population. We randomly equalised the sample size of each population to that of the smallest (n=7). The filtered SNPs from the isolation-by-distance analysis above were combined with invariant sites, and site-level information was extracted for the selected individuals from the downsampling. Mean **π** and Tajima’s D were calculated for 500 kbp non-overlapping windows across the genome using the software pixy v. 2.0.0 (Korunes and Samuk, 2021), windows without RAD-loci were discarded. The genome-wide weighted mean of **π** was calculated for each population, as recommended by Samuk (2022). To test whether populations differed with respect to each statistic, a Kruskal-Wallis test was performed, and to identify pairwise differences between populations, a post-hoc Dunn test was used with Benjamini-Hochberg FDR correction.

Inbreeding coefficients (F_IS_) were calculated for the same subsetted (n=7) populations generated above. ANGSD (with the parameters: -doHWE 1 -domajorminor 1 -GL 1 -doMaf 1 -SNP_pval 1e-4 -minInd 7) generated genotype likelihoods and performed Hardy-Weinberg equilibrium tests to calculate F_IS_ for each variable site across all individuals per population. A mean F_IS_ was then generated for each population.

### Genome scan for local adaptation

A PCA-based genome scan for signatures of local adaptation was carried out on all the Northern European populations using the R package pcadapt v.4.4.1(Privé et al. 2020). Populations from Portugal and Studland (UK) were excluded as these were highly differentiated and dominated the PCAs (see results). By removing them we were able to focus on a northeastern gradient along the coast from Belgium to Sweden to identify potential loci associated with local adaptation. An optimal number of genetic clusters (*k*) was determined based on a scree plot of the variance explained by each component. A Mahalanobis distance test was used to calculate correlations between variants and principal components. Significance was determined using a false discovery rate (FDR) adjusted q-value of <0.05.

To infer whether the outlying SNPs (FDR-adjusted q-value < 0.05) were associated with specific biological functions we performed a Gene Ontology (GO) enrichment analysis. First, *C. arenaria* genes were functionally annotated by identifying homologous sequences in *Arabidopsis thaliana* (taxonomy ID: 3702) and *Oryza sativa* (taxonomy ID: 4530) proteomes from UniProt, and retaining their associated GO terms. Homologous sequences were identified using blastp with e-value cutoff of 1e-5, retaining only the top 5 matches. GO enrichment was then assessed for genes located within 50 kb windows surrounding each outlying SNP (‘outlier genes’ hereafter), using all retained SNPs as the background set. The enrichment analysis was performed using the weightCount algorithm from the R package topGO v2.58 (Alexa and Rahnenführer, 2017), which uses Fisher’s exact test to assess enrichment while accounting for dependencies between related GO terms. GO terms with less than three significant genes in the test set were discarded. Protein and gene name information was retrieved from the Uniprot database for outlier genes with Uniprot matches.

## Results

### A chromosome scale reference genome for marram grass

The flow cytometry inferred 1C genome size for the UK-SFB-005 accession was 4.37 Gb. From this accession we generated 277.29 Gb of PacBio HiFi data with an N50 read length of 19.1 kb, and a coverage of 73x. The genome scope plot of the HiFi reads displays four peaks in the kmer spectra (Figure S1), indicating UK-SFB-005 is likely a tetraploid. This matches with our expectation, as ∼90% of the 36 *C. arenaria* accessions in the Chromosome Count Database (Rice et al., 2014) are tetraploid (2n = 4*x* = 24), with only a single diploid (2n = 2x = 14) and octoploid (2n = 8x = 48) reported in the 1940s and 1950s, respectively, with each appearing twice in the database despite originating from a single source. Our flow cytometry estimate is also very similar to the 1C value of 4.29 Gb that was estimated for a tetraploid individual with 2n = 28 (Tkach *et al.,* 2025). The hifiasm genome assembly was 3.8 Gb in length, highly contiguous (N90 = 103.25 Mb; L90 = 13), and complete (BUSCOs 99.3% complete, 85.8% duplicated, 0.1% fragmented). The Helixer annotation identified 107,132 protein coding genes, with the same percentage of complete and duplicated BUSCOs as the assembly itself. The separate assembly of both subgenomes allowed us to map ddRAD-seq reads to their respective subgenomes and perform downstream SNP calling under a diploid model.

### Gene family expansions associated with adaptation to the coastal environment

The OrthoVenn3 analysis identified 40,529 orthologous gene clusters, of which 10,835 were shared across all five included species (Figure S2). Relative to the other taxa included in the analysis, *C. arenaria* showed significant enrichment of 199 GO terms among expanded gene families and 49 GO terms among contracted gene families (Figure S3; Tables S2 and S3). Several enriched terms were associated with biological processes relevant to the environmental conditions characteristic of coastal dunes.

Expanded gene families showed strong enrichment for functions associated with abiotic stress, particularly osmotic, ionic and water stress (Figure 1). Response to osmotic stress was significantly enriched among expanded families (8 families, *p* < 0.001), as was response to salt stress (23 families, *p* < 0.001). Response to water deprivation was also enriched (154 families, *p* < 0.001), together with response to abscisic acid (24 families, *p* < 0.001) and stomatal closure (10 families, *p* < 0.001). Gene families associated with ionic regulation were also overrepresented, alongside strong enrichment of the response to oxidative stress (118 gene families, *p* < 0.001)

**Figure 1:**
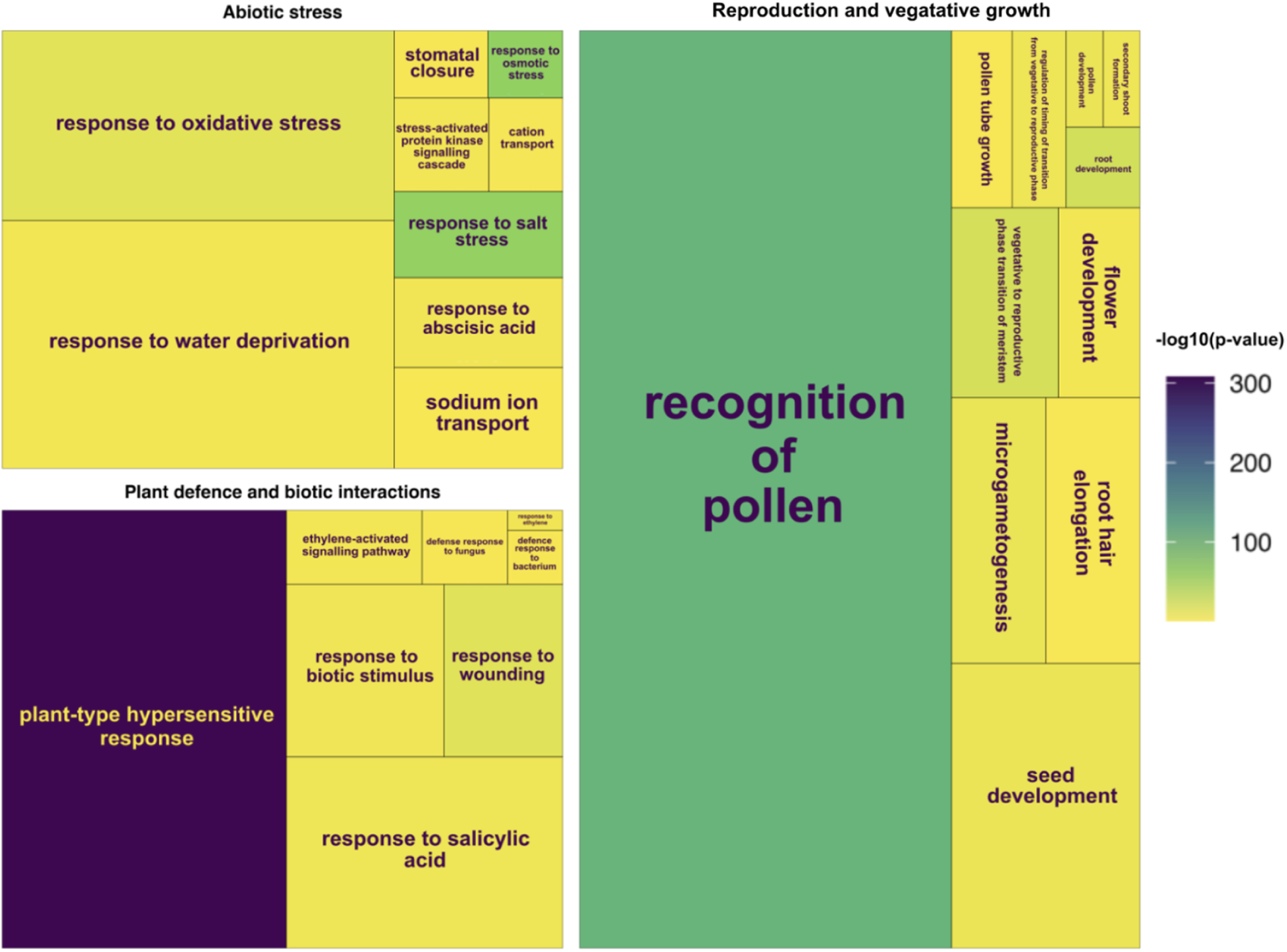
Treemap of significantly enriched Gene Ontology (GO) terms associated with abiotic stress tolerance, reproduction and plant defence in gene families expanded in *C. arenaria* relative to the other species included in the CAFE5 analysis. Block size represents the number of gene families associated with each GO term, while colour indicates the significance of enrichment based on Fisher’s exact tests (see key). All GO terms shown are significantly enriched (*p* < 0.05). Full results are provided in Table S2.

Expanded families were also enriched for functions associated with biotic stress and plant development. Response to biotic stimulus was represented by 73 expanded families (*p* < 0.001), with additional enrichment for response to salicylic acid (142 families, *p* < 0.001), ethylene-activated signalling (27 families, *p* < 0.001), and defence responses to fungi (17 families, *p* < 0.001). Developmental processes associated with reproduction and below ground and vegetative growth were also enriched, including recognition of pollen (286 families, *p* < 0.001), root development (5 families, *p* < 0.001), root hair elongation (21 families, *p* < 0.001) and secondary shoot formation (3 families, *p* < 0.001).

### Evident population structure within Europe

In total, we generated sequence data for 266 *C. arenaria* samples from the 10 populations (Figure 2A), with 239 after removing samples with insufficient data quantity or quality (mean number of samples per population = 24; SD = 13; range = 7-45; Table S1) Across the whole sampling range, the largest component of genetic variation was explained by differences between the Portuguese populations and the rest of the samples from Northwestern Europe (31.8% of total variation, Figure 2B; F_ST_ mean = 0.602; SD = 0.017). To look at the finer population structure within the Northwestern European populations, the Portuguese populations were removed from the dataset, which revealed clustering largely explained by geography. The largest component of variation of this reduced dataset, comprising 222 samples, split the population from Studland on the south coast of the UK (‘UK_stu’) from the rest of the populations (3.32% of total variation, Figure 2C). For the Northwestern European populations, admixture analysis indicated *K* = 2 as the most likely number of populations (Figure S4). The admixture assignments of each individual in the dataset mirrored the clustering seen in the PCA (Figure 2D). A northeastern cline is apparent in both the PCA and admixture results, with the notable exception of Sweden which shows a within-site split, with one distinct group showing no admixture (‘SE_yst1’), and one group showing greater similarity with North Sea populations (‘SE_yst2’) (Figure 2D).

**Figure 2.**
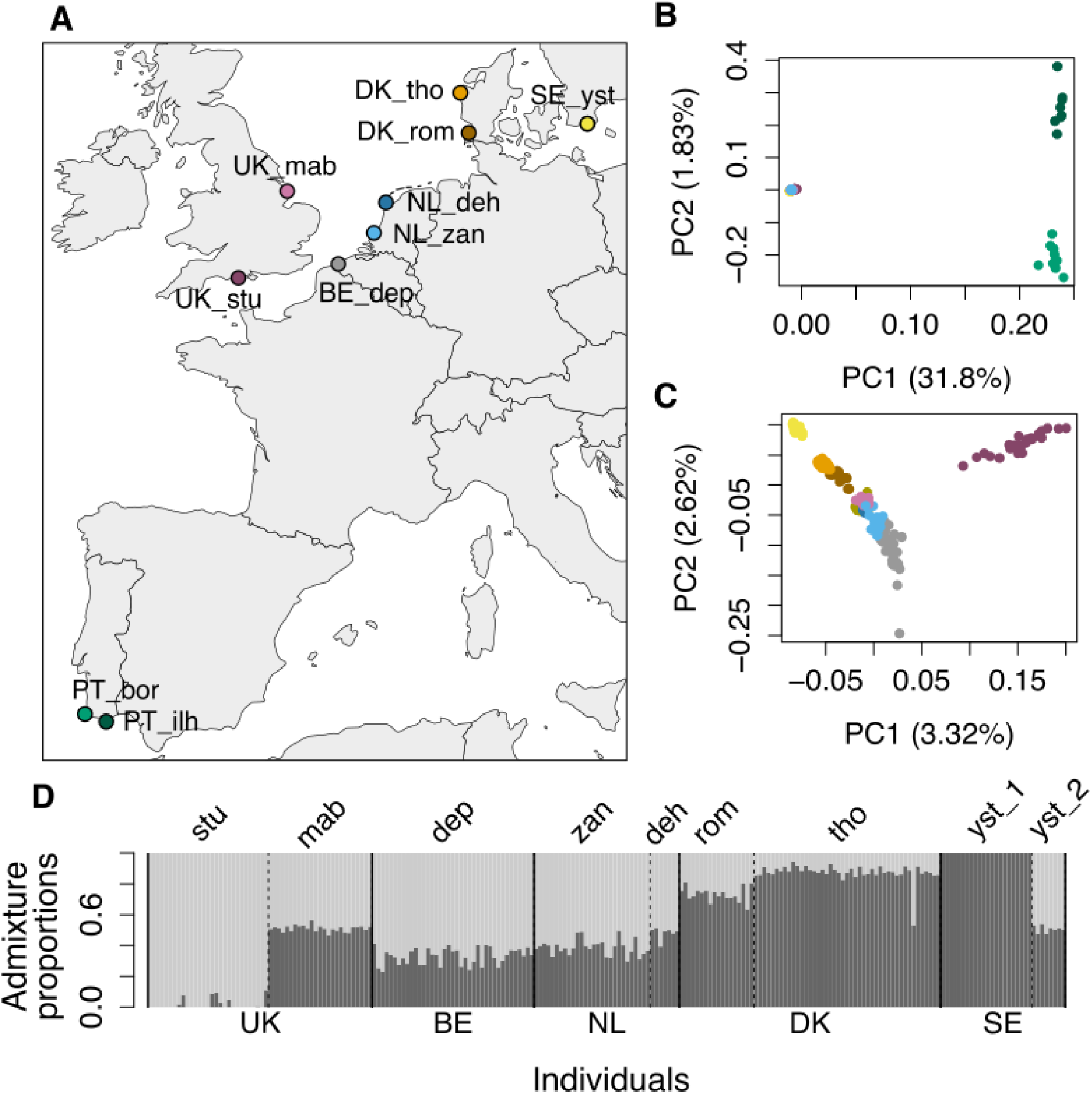
Population structure of *Calamagrostis arenaria* across its native European range. **A.** Map of populations of *C. arenaria* that were sampled and genotyped with RAD sequencing. Colours of points on the map continue to indicate populations in subsequent plots. **B.** PCA of all sampled populations. **C.** PCA excluding samples from Portugal revealing finer structure in Northwestern Europe (darker yellow points indicate individuals from the ‘SE_yst2’ group). **D.** Estimates of admixture proportions at *K=2*. Country codes are shown below the admixture proportions, and countries are separated by solid lines. Population names are shown above the admixture proportions, if there are multiple populations per country these are separated by dashed lines.

A maximum likelihood tree based on 100,844 SNPs supports the patterns observed in the PCA and admixture analyses (Figure 3A & S5). The tree has a deep split between the Portuguese and Northwestern European samples. The Swedish population is again divided into two groups, with individuals from ‘SE_yst2’ clustering largely with Dutch populations (Figure 3A).

**Figure 3.**
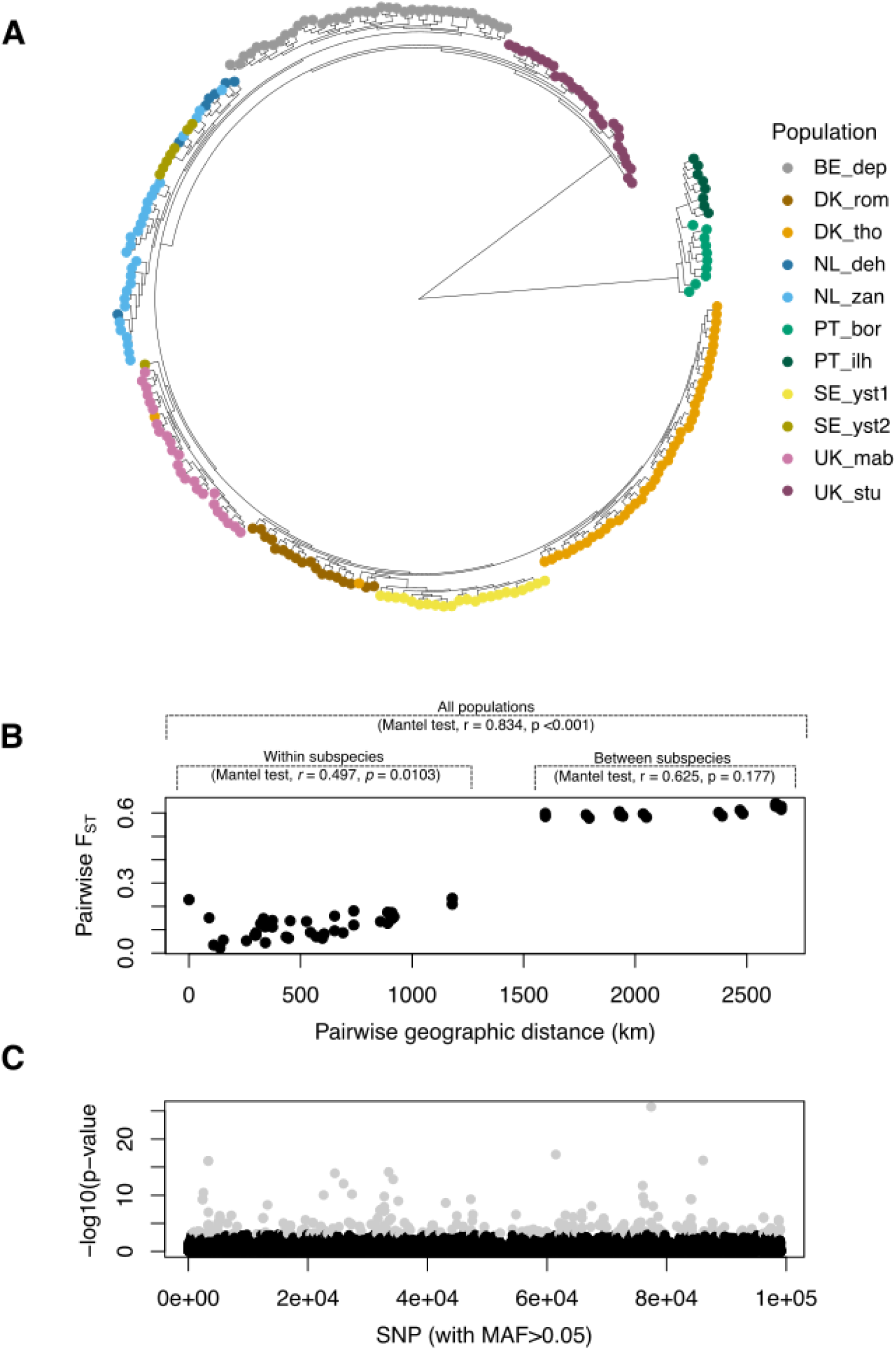
**A.** Phylogenetic relationships amongst individuals. The native individuals from Ystad, Sweden (‘SE_yst_1’) and the introduced individuals to the same beach (‘SE_yst_2’) are indicated on the tree (a linear version of the phylogenetic tree, together with additional individual level information is given in Figure S5. **B.** The relationship between pairwise genetic distance (F_ST_) and pairwise geographic distance (km). **C.** SNPs showing a significant association with population structure (light grey).

### Isolation-by-distance within subspecies but not between

There is a strong positive correlation between geographic and genetic distance across all pairwise population comparisons (Mantel test, *r* = 0.834, *p* <0.001) (Figure 3B). Between the Northwestern European populations of *C. arenaria* subsp. *arenaria* there is also a positive correlation (Mantel test, *r* = 0.497, *p* = 0.0103), but between Portuguese (*C. arenaria* subsp. *australis*) and Northwestern European populations there is no significant pattern of isolation-by-distance (Mantel test, *r* = 0.625, *p* = 0.177).

### Genome-wide diversity and signatures of demography and inbreeding

Genome-wide mean genetic diversity (**π**) calculated for 500 kb windows of the genome was generally low (mean = 0.00107, SD = 0.00163, Figure S6A), although it did differ significantly among populations (Kruskal-Wallis rank sum test, ◻^2^= 340.87, d.f. = 10, *P* < 0.001). A post-hoc Dunn’s test revealed significant pairwise differences between populations (Table S4). An independently calculated weighted population average of **π** for each population showed the same pattern, with highest genetic diversity in the Portuguese populations (PT_ilh and PT_bor), and lowest in the subdivided Swedish population (SE_yst1 and SE_yst2) (Table 1). Tajima’s D was moderately positive across all populations (mean = 0.955; SD = 0.481, Figure S6B) which can be due to recent population contractions. F_IS_ inbreeding coefficients for each population ranged from slightly negative to slightly positive (-0.159 – 0.068, mean = -0.002; SD = 0.056). Most populations had a positive F_IS_ (Table 1), which indicates a deficit of heterozygotes, which can result from inbreeding, but could also be explained by clonal reproduction (Stoeckel et al. 2006). SE_yst1, SE_yst2, and UK_stu all had a slightly negative F_IS_, indicating an excess of heterozygotes which can result from outcrossing.

**Table 1:**
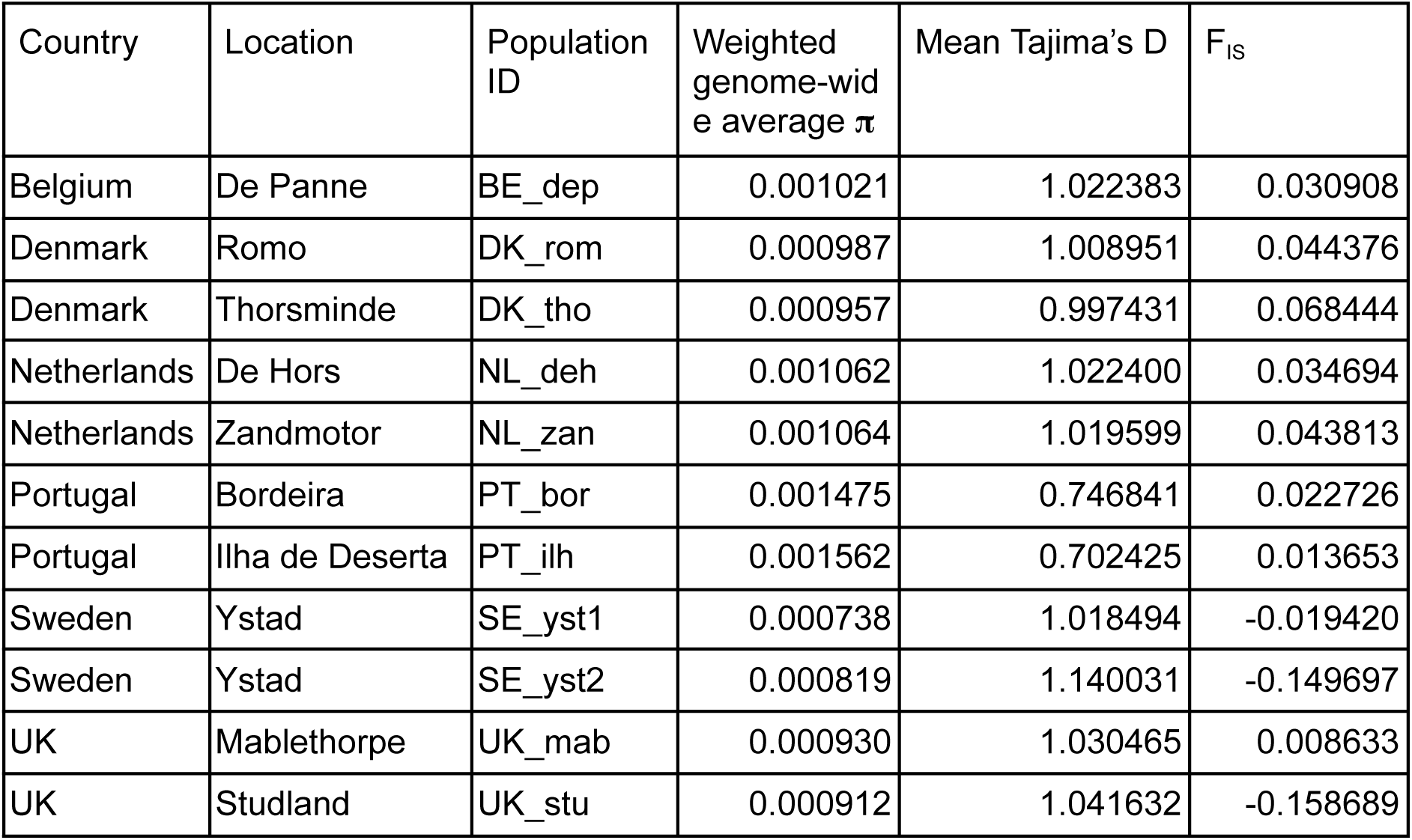
Genetic diversity summary statistics per population.

| Country | Location | Population ID | Weighted genome-wide average **π** | Mean Tajima’s D | F<sub>IS</sub> |
|---|---|---|---|---|---|
| Belgium | De Panne | BE_dep | 0.001021 | 1.022383 | 0.030908 |
| Denmark | Romo | DK_rom | 0.000987 | 1.008951 | 0.044376 |
| Denmark | Thorsminde | DK_tho | 0.000957 | 0.997431 | 0.068444 |
| Netherlands | De Hors | NL_deh | 0.001062 | 1.022400 | 0.034694 |
| Netherlands | Zandmotor | NL_zan | 0.001064 | 1.019599 | 0.043813 |
| Portugal | Bordeira | PT_bor | 0.001475 | 0.746841 | 0.022726 |
| Portugal | Ilha de Deserta | PT_ilh | 0.001562 | 0.702425 | 0.013653 |
| Sweden | Ystad | SE_yst1 | 0.000738 | 1.018494 | -0.019420 |
| Sweden | Ystad | SE_yst2 | 0.000819 | 1.140031 | -0.149697 |
| UK | Mablethorpe | UK_mab | 0.000930 | 1.030465 | 0.008633 |
| UK | Studland | UK_stu | 0.000912 | 1.041632 | -0.158689 |

### Genome scan for local adaptation identifies stress response loci

We scanned for genomic signatures of local adaptation by identifying SNPs associated with population structure. According to a scree plot (Figure S7), *k*=7 optimally described population structure. We detected 334 out of 99,076 SNPs (0.337%) as significant outliers (*q* < 0.05; Figure 3C). 858 genes were within 50 kb of the significantly associated SNPs, and were enriched for 10 GO terms (Table S5). Of the candidate genes, the six biological process-enriched GO-terms were largely associated with responses to nutrient limitation, abiotic stress, and pathogens (Table S5). Of the 858 candidate genes, 445 were successfully functionally annotated from the Uniprot database (Table S6), identifying a range of functions associated with nutrient deficiency, drought and salt tolerance, pathogen defence, lignin biosynthesis and mechanical strength.

## Discussion

*Calamagrostis arenaria* plays a crucial role in forming the sand dunes that protect a vast stretch of coastlines around Europe from erosion, flooding and extreme weather events. Here we provide the first genome-level insights into how natural processes and human intervention have shaped genetic variation in the species.

### Expansion of gene families associated with the coastal environment

The enrichment of expanded gene families associated with salt stress (GO:0009651), osmotic stress (GO:0006970), water deprivation (GO:0009414) and oxidative stress (GO:0006979) suggests that abiotic stress responses are an important component of the *C. arenaria* genomic repertoire. These functions are consistent with the environmental pressures of coastal dunes, where salt spray, rapidly draining sandy substrates and fluctuating water availability challenge water and ion homeostasis (Du and Hesp, 2020; Hesp and Martínez, 2007; Maun, 2009; Whitford, 2002). Enrichment of abscisic acid response (GO:0009737) and stomatal closure (GO:0090332) further implicates pathways regulating water loss, while sodium ion transport (GO:0006814) may contribute to ionic homeostasis under saline conditions. Together, these patterns suggest that gene-family expansion in *C. arenaria* involves multiple interconnected components of abiotic stress responses.

Expanded gene families were also enriched for root development (GO:0048364), root hair elongation (GO:0048767) and secondary shoot formation (GO:0010223). These functions may support the extensive root systems and clonal propagation that enable *C. arenaria* to thrive as sand accumulates (Maun, 1998, 2009; van der Lan et al., 1997). Alongside enriched defence-related processes, these expansions may contribute to the coordination of vegetative growth and biotic interactions in dynamic dune environments (Koske et al., 2004). As gene-family expansions and contractions were inferred relative to *Lolium perenne*, these changes could have occurred at any point since the divergence of the two lineages, encompassing multiple speciation events, rather than necessarily being specific to *C. arenaria*. Comparisons with additional, more closely related species would help resolve when these changes occurred, while including other dune-building grasses would establish whether similar patterns have evolved convergently in other similarly adapted species.

### North–south genetic divergence in European marram grass

We find strong genetic differentiation between southern and northern European populations (Figure 2B, 3A & 3B). This reflects the two recognised European subspecies of marram grass, with one found in northern and western Europe (*C. arenaria* subsp. *arenaria*), and the other in southern and mediterranean regions (*C. arenaria* subsp. *australis*). The division between marram from these two regions has been recognised for over a hundred years (Ascherson & Graebner, 1899), and they can be morphologically distinguished using differences in the structure of glumes and hairs at the base of the spikelets. Previous research using inter-simple sequence repeat (ISSR) markers also reported strong differentiation between subspecies, with samples from northern and southern Atlantic coast of Portugal and the French Mediterranean coast (Camargue) being distinct from northern European individuals (Rodríguez-Echeverría et al. 2008).

Given the strong genetic differentiation between subspecies (F_ST_ mean = 0.602; SD = 0.017), together with diagnostically reliable morphological differences, these lineages may be better interpreted as separate species. However, the extent of gene flow between subspecies remains hard to ascertain from our results alone as we lack samples from locations near their range overlap on the French Basque Coast (Figure 3B). It is important to note that this ‘distance gap’ is partially an artefact of using euclidean distances in our analysis. When using more ecologically realistic routes (i.e. measuring minimum distances by sea), the geographic distance between northern populations where we have gene flow (e.g. Studland, UK to Ystad, Sweden) are analogous with the distances between subspecies (e.g. Studland, UK to Bordeira, Portugal), both being ∼1,700 Km.

Overall, between subspecies we did not detect a significant positive correlation between geographic and genetic distances. However, distinguishing isolation-by-distance from reproductive isolation will require denser sampling along the Western European Atlantic coast. Such sampling would clarify whether the subspecies form a continuous distribution with a potential hybrid zone, or are instead separated by breaks in suitable dune habitat such as the rocky coastline of northern Spain.

### Coastal genetic structure and historical vicariance in northern C. arenaria

Within the northern *C. arenaria* subsp. *arenaria,* genetic structure largely reflects geographical distribution, with a strong pattern of isolation by distance among populations (Figure 3B) consistent with ongoing gene flow between neighbouring populations. An exception to this pattern is the southern UK population from Studland, which is strongly differentiated from all other populations, including the eastern UK population from Mablethorpe (Figure S5) which clusters with the Dutch populations despite being separated by ∼300 km of the North Sea (Figure 2A). Although connectivity across open water is likely to be readily feasible through the rafting of rhizomes which can remain viable for up to 70 days in seawater (Konlechner and Hilton 2009).

A similar disjunct generic structure along the UK coastline was reported using ISSR markers (Rodríguez-Echeverría et al. 2008), where populations from the west coast of the UK (Ynyslas, Wales) were genetically distinct from the east coast (Blakeney Point, Norfolk) and mainland Europe. A plausible explanation for this pattern is the legacy of quaternary glaciations and associated sea-level fluctuations which repeatedly sealed the English channel, dramatically decreasing the connectivity of UK populations either side of this land bridge, although further sampling is required to confirm this. The changing patterns of ice coverage and sea levels due to glaciation has had a lasting impact on the genetic diversity of the flora and fauna of the British Isles (Fenton et al. 2023), and glacial history is one of the major drivers of genetic discontinuities in coastal marine species around northwest Europe (Pagowski & Micheli, 2024).

### Signatures of local adaptation across northern dune populations

Genetic variants that significantly contributed to population structure within the northern subspecies *C. arenaria* subsp. *arenaria* were associated with genes that were enriched for biological processes related to a range of environmental stressors relevant to the coastal environment.

Dunes are formed due to a dynamic feedback loop whereby marram grass captures sand thereby driving dune formation, while sand accumulation, in return, impacts marram performance and survival (Bonte et al. 2021). Young plants must be able to withstand burial and wind exposure to establish, and we find evidence of potential local adaptation at genes involved in lignin biosynthesis and mechanical strength. Genes encoding Cinnamyl alcohol dehydrogenase 7 (alternative name Flexible culm 1 [FC1]), and Kinesin-like protein KIN-4A (alternative name Brittle culm 2 [BC2]) which are described in rice (Li et al. 2009; Zhang et al. 2010), may influence the balance between mechanical strength and flexibility required to withstand sand accumulation and wind exposure in marram. Lack of burial, however, is also a threat to marram, as regular sand burial provides marram grass with the opportunity to grow roots in wind-blown sediment free of root-feeding nematodes or pathogenic fungi (Van der Putten et al. 1988, 1989). There is significant enrichment of the biological processes ‘response to nematode’ (GO: 0031969) and ‘defence response to fungus’ (GO:0050832), potentially reflecting spatial variation in pathogen-mediated selection. Previous studies have demonstrated that soil-borne pathogens influence marram grass performance, and that nematode and fungal communities differ between dune systems, including those with contrasting soil chemistry (De Rooij-van der Goes et al., 1995; Van De Walle et al., 2023). Together, these findings suggest that local adaptation shapes the feedback between marram growth and sand accumulation, influencing both the capacity of plants to build dunes through sand capture and to persist under contrasting burial regimes, including periods of low burial when exposure to soil-borne antagonists may increase.

Exposure to drying winds, salt spray and nutrient-deficient substrates also varies substantially among geographically distinct coastlines, and several of the outlying regions may contribute to the mitigation of these stressors. For example, phosphatidylinositol phosphate biosynthetic process (GO:0046854) is enriched among outlying SNPs (Figure 3C). This pathway contributes to the production and turnover of phosphoinositides, lipid signalling molecules that play key roles in regulating plant drought and salt stress responses (Munnik & Vermeer, 2010). PIP5K9 is a candidate gene linked to this GO term, which impacts root growth under salt stress conditions (Kuroda et al. 2021). Other candidate genes such as MYB52, connected with drought, salt tolerance and ion homeostasis (Park et al. 2011; Di et al. 2026), and Salt-Related MYB 1 (SRM1), involved in salt stress tolerance (Wang et al. 2015), suggest local adaptation in response to differences in salinity. Coastal dunes are nutrient poor (Willis and Yemm 1961), and outlying genes include those associated with methionine metabolic process (GO:0006555). Methionine is a precursor for S-adenosylmethionine (SAM), levels of which can signal stress and nutrient limitations (Fang et al. 2023).

Adaptation to different climatic conditions across the sampling range is likely, as the southernmost population in this analysis (De Panne, Belgium) will experience higher annual temperatures (+1.44°C) and lower total rainfall (-12%) compared to the northernmost populations (Thorsminde, Denmark). Signatures of local adaptation were detected in genes connected to heat and drought tolerance: SUD1, mutations of which have been shown to suppress drought-sensitive phenotypes in *A. thaliana* (Doblas et al. 2013); and *HSP83*, which is part of a family of heat shock proteins that show upregulation in response to heat and play a role in plant defences against high temperatures (Kushawaha et al. 2021).

Overall, the identification of genetic variation associated with population structure provides evidence for local adaptation to a diverse range of coastal stressors. These findings highlight the importance of incorporating local provenance into dune restoration strategies to maximise restoration success while minimising the potential disruption or loss of locally adapted genetic variation.

### Genetic shadow of dune restoration but no obvious gene flow

The Ystad population in Sweden comprised two distinct genetic clusters: one genetically unique group (SE_yst1), likely representing native Swedish plants at the northern edge of the species’ genetic cline, and a second group (SE_yst2) clustering with Dutch populations alongside a single individual grouping with UK samples (Figures 2 & 3). As this site has undergone dune restoration involving *C. arenaria* replanting (most recently in 2020), these patterns suggest that the introduced plants were likely sourced from outside the local population, and likely from The Netherlands.

Both Swedish clusters exhibited low genetic diversity and slightly negative inbreeding coefficients, consistent with high clonality. Positive Tajima’s D values further suggest recent population contraction, which may explain the need for restoration. The lack of strong admixture between native and introduced plants indicates limited gene flow so far, although the short period (one year) between the latest restoration work and sampling for this work may limit detection of introgression. Introducing genetically depauperate material can reduce individual fitness and constrain population growth (Williams 2001), as observed in dune restoration of *Calamagrostis breviligulata* in USA, where a monotypic commercial variety reduced genetic diversity (Slaymaker et al., 2015). In Ystad, replanting has increased genetic diversity in the short term, but the long-term outcome remains uncertain, including whether hybrids will suffer from outbreeding depression, whether a single clone will dominate, or whether diversity will be enhanced over longertime scales. Given the widespread evidence of local adaptation, the use of non-local material may ultimately compromise restoration success, as locally sourced material is likely to show greater persistence.

## Conclusions

Our chromosome-scale genome and population genomic analyses shed new light on the evolutionary and ecological processes that have shaped *Calamagrostis arenaria* as a dominant coastal dune builder. We identify genomic signatures of adaptation to the challenging dune environment, strong lineage divergence across Europe, and genetic consequences of restoration using non-local material. Together, these findings highlight the importance of conserving locally adapted genetic variation and integrating genomic approaches into future dune restoration and coastal conservation strategies.

## Acknowledgements

EC, EJB & CL are supported by a Human Frontiers Science Project (HFSP) research grant (RGEC28/2024); ERB is supported by a Natural Environment Research Council (NERC) Adapting to the Challenges of a Changing Environment (ACCE) studentship (NE/S00713X/1); the reference genome was assembled with funding through a NERC Environmental Omics Facility (NEOF) de novo genome pilot competition (NEOF1365); LTD was supported by a NERC Independent Research Fellowship (NE/T011025/1). VCR was supported by NWO-Veni grant VI.Veni.212.059.

## Data Accessibility and Benefit-Sharing

All raw whole-genome sequencing, ddRAD-seq data, and the UK-SFB-005 reference genome and annotation have been deposited with NCBI under the Bioproject PRJNA1543376. All scripts used in this study are available on GitHub: https://github.com/Sheffield-Plant-Evolutionary-Genomics/

## Author Contributions

LJ, NJN, VCR & LTD conceived and designed the research with contributions from all coauthors. CL, SGSH, ERB, MV, VCR, EW, JG & LTD conducted fieldwork and identified specimens. EJB, ERB, RP, LSK, SM, IJL conducted lab work. ERB, EVC, LTD and NN analysed and interpreted the population and genomic data. ERB, EVC & LTD wrote the manuscript with input from all authors.

## Notes

### Competing Interest Statement

The authors have declared no competing interest.

## References

Alexa, A. and Rahnenführer, J. (2017) topGO: Enrichment analysis for Gene Ontology. Bioconductor. doi:10.18129/B9.BIOC.TOPGO.

Bansal, J., Gupta, K., Rajkumar, M.S., Garg, R. and Jain, M. (2021) ‘Draft genome and transcriptome analyses of halophyte rice Oryza coarctata provide resources for salinity and submergence stress response factors’, Physiologia Plantarum, 173(4), pp. 1309–1322. doi:10.1111/ppl.13284.

Bar, P., Katz, O. and Dorman, M. (2023) ‘Spatial Heterogeneity Effects on Meta-Community Stability of Annual Plants from a Coastal Dune Ecosystem’, Plants, 12(11), p. 2151. doi:10.3390/plants12112151

Bogdanović, S., Šegota, V. and Alegro, A. (2018) ‘Resurrection of the regionally extinct taxon in Croatia – the case of Ammophila arenaria (L.) Link (Poaceae)’, Acta Botanica Croatica, 77(2), pp. 214–217. doi:10.2478/botcro-2018-0013.

Bonte, D., Batsleer, F., Provoost, S., Reijers, V., Vandegehuchte, M.L., Van De Walle, R., Dan, S., Matheve, H., Rauwoens, P., Strypsteen, G., Suzuki, T., Verwaest, T. and Hillaert, J. (2021) ‘Biomorphogenic feedbacks and the spatial organization of a dominant grass steer dune development’, Frontiers in Ecology and Evolution, 9, 761336. doi:10.3389/fevo.2021.761336.

Borsje, B.W., van Wesenbeeck, B.K., Dekker, F., Paalvast, P., Bouma, T.J., van Katwijk, M.M. and de Vries, M.B. (2011) ‘How ecological engineering can serve in coastal protection’, Ecological Engineering, 37(2), pp. 113–122. doi:10.1016/j.ecoleng.2010.11.027.

Breed, M.F., Stead, M., Ottewell, K., Gardner, M. and Lowe, A. (2013) ‘Which provenance and where? Seed sourcing strategies for revegetation in a changing environment’, Conservation Genetics, 14(1), pp. 1–10. doi:10.1007/s10592-012-0425-z.

Chang, C.C., Chow, C.C., Tellier, L.C.A.M., Vattikuti, S., Purcell, S.M. and Lee, J.J. (2015) ‘Second-generation PLINK: rising to a new standard of genetic analysis’, GigaScience, 4, 7. doi:10.1186/s13742-015-0047-8.

Chen, S., Zhou, Y., Chen, Y. and Gu, J. (2018) ‘fastp: an ultra-fast all-in-one FASTQ preprocessor’, Bioinformatics, 34(17), pp. i884–i890. doi:10.1093/bioinformatics/bty560.

Cheng, H., Concepcion, G.T., Feng, X., Zhang, H. and Li, H. (2021) ‘Haplotype-resolved de novo assembly using phased assembly graphs with hifiasm’, Nature Methods, 18, pp. 170–175.

Clark, J., Hidalgo, O., Pellicer, J., Liu, H., Marquardt, J., Robert, Y., Christenhusz, M. et al. (2016) ‘Genome evolution of ferns: evidence for relative stasis of genome size across the fern phylogeny’, New Phytologist, 210, pp. 1072–1082.

Cullingham, C.I., Miller, J.M., Peery, R.M., Dupuis, J.R., Malenfant, R.M., Gorrell, J.C. and Janes, J.K. (2020) ‘Confidently identifying the correct K value using the ΔK method: When does K = 2?’, Molecular Ecology, 29(5), pp. 862–869.

De Lillis, M., Costanzo, L., Bianco, P.M. and Tinelli, A. (2004) ‘Sustainability of sand dune restoration along the coast of the Tyrrhenian Sea’, Journal of Coastal Conservation, 10(1), pp. 93–100. 10.1007/BF02818945

De Rooij-van der Goes, P.C.E.M. (1995) ‘The role of plant-parasitic nematodes and soil-borne fungi in the decline of Ammophila arenaria (L.) Link’, New Phytologist, 129(4), pp. 661–669. 10.1111/j.1469-8137.1995.tb03035.x

Devos, K.M. et al. (2023) ‘Genome analyses reveal population structure and a purple stigma color gene candidate in finger millet’, Nature Communications, 14, article 3694. 10.1038/s41467-023-38915-6

Di, Y., Lou, S., Wang, Z., Wang, T., Niu, X., Ji, Z., Liu, W., Chen, S., Guo, Z., Zheng, S., Zhang, T., Piao, F., Wang, Y., Dong, X., Hu, C. and Dong, H. (2026) ‘The transcription factor MYB52 regulates salt stress tolerance in tomato by modulating ion homeostasis and proline biosynthesis’, Plant Physiology, 201(1), article kiag256. doi:10.1093/plphys/kiag256.

Doblas, V.G., Amorim-Silva, V., Posé, D., Rosado, A., Esteban, A., Arró, M., Azevedo, H., Bombarely, A., Borsani, O., Valpuesta, V., Ferrer, A., Tavares, R.M. and Botella, M.A. (2013) ‘The SUD1 gene encodes a putative E3 ubiquitin ligase and is a positive regulator of 3-hydroxy-3-methylglutaryl coenzyme A reductase activity in Arabidopsis’, The Plant Cell, 25(2), pp. 728–743. doi:10.1105/tpc.112.108696.

Doležel, J., Greilhuber, J. and Suda, J. (2007) ‘Estimation of nuclear DNA content in plants using flow cytometry’, Nature Protocols, 2, pp. 2233–2244.

Du, J. and Hesp, P.A. (2020) ‘Salt spray distribution and its impact on vegetation zonation on coastal dunes: a review’, Estuaries and Coasts, 43(8), pp. 1885–1907. doi:10.1007/s12237-020-00820-2.

Evanno, G., Regnaut, S. and Goudet, J. (2005) ‘Detecting the number of clusters of individuals using the software STRUCTURE: a simulation study’, Molecular Ecology, 14(8), pp. 2611–2620.

Fang, W., Jiang, L., Zhu, Y., et al. (2023) ‘Methionine restriction constrains lipoylation and activates mitochondria for nitrogenic synthesis of amino acids’, Nature Communications, 14, article 2504. doi:10.1038/s41467-023-38289-9.

Fenton, S., Elmer, K.R., Bean, C.W. and Adams, C.E. (2023) ‘How glaciation impacted evolutionary history and contemporary genetic diversity of flora and fauna in the British Isles’, Scottish Geographical Journal, 139(3–4), pp. 445–465. doi:10.1080/14702541.2023.2231407.

Fernández-Montblanc, T., Duo, E. and Ciavola, P. (2020) ‘Dune reconstruction and revegetation as a potential measure to decrease coastal erosion and flooding under extreme storm conditions’, Ocean & Coastal Management, 188, 105075. doi:10.1016/j.ocecoaman.2019.105075.

Gao, J., Kennedy, D.M. and Konlechner, T.M. (2020) ‘Coastal dune mobility over the past century: a global review’, Progress in Physical Geography: Earth and Environment, 44(6), pp. 814–836. doi:10.1177/0309133320919612.

Gilbert, M.E., Pammenter, N.W. and Ripley, B.S. (2008) ‘The growth responses of coastal dune species are determined by nutrient limitation and sand burial’, Oecologia, 156(1), pp. 169–178. doi:10.1007/s00442-008-0968-3.

Hesp, P.A. (1991) ‘Ecological processes and plant adaptation on coastal dunes’, Journal of Arid Environments, 21, pp. 165–191.

Hesp, P.A. (2002) ‘Foredunes and blowouts: initiation, geomorphology and dynamics’, Geomorphology, 48, pp. 245–268.

Hesp, P.A. and Martínez, M.L. (2007) ‘Disturbance processes and dynamics in coastal dunes’, in Johnson, E.A. and Miyanishi, K. (eds.) Plant Disturbance Ecology. San Diego: Academic Press, pp. 215–247.

Hijmans, R.J. (2025) geosphere: Spherical Trigonometry. R package version 1.5-20. Available at: CRAN. doi:10.32614/CRAN.package.geosphere.

Hoffmann, A.A., Miller, A.D. and Weeks, A.R. (2021) ‘Genetic mixing for population management: From genetic rescue to provenancing’, Evolutionary Applications, 14(3), pp. 634–652. doi:10.1111/eva.13154.

Hol, W.H.G., van der Wurff, A.W.G., Skøt, L. and Cook, R. (2008) ‘Two distinct AFLP types in three populations of marram grass (Ammophila arenaria) in Wales’, Plant Genetic Resources: Characterization and Utilization, 6(3), pp. 201–207. doi:10.1017/S147926210899417X.

Holst, F., Bolger, A.M., Kindel, F. et al. (2025) ‘Helixer: Ab initio prediction of primary eukaryotic gene models combining deep learning and a hidden Markov model’, Nature Methods. doi:10.1038/s41592-025-02939-1.

Huiskes, A.H.L. (1977) ‘The natural establishment of Ammophila arenaria from seed’, Oikos, 29, pp. 133–136

Konlechner, T.M. and Hilton, M.J. (2009) ‘The potential for marine dispersal of Ammophila arenaria (marram grass) rhizome in New Zealand’, Journal of Coastal Research, pp. 434–437.

Kopelman, N.M., Mayzel, J., Jakobsson, M., Rosenberg, N.A. and Mayrose, I. (2015) ‘CLUMPAK: a program for identifying clustering modes and packaging population structure inferences across K’, Molecular Ecology Resources, 15(5), pp. 1179–1191. doi:10.1111/1755-0998.12387.

Korneliussen, T.S., Albrechtsen, A. and Nielsen, R. (2014) ‘ANGSD: Analysis of Next Generation Sequencing Data’, BMC Bioinformatics, 15, 356. doi:10.1186/s12859-014-0356-4.

Korunes, K.L. and Samuk, K. (2021) ‘pixy: Unbiased estimation of nucleotide diversity and divergence in the presence of missing data’, Molecular Ecology Resources, 21(4), pp. 1359–1368. doi:10.1111/1755-0998.13326.

Koske, R.E., Gemma, J.N., Corkidi, L., Siguenza, C. and Rincon, E. (2004) ‘Arbuscular mycorrhizas in coastal dunes’, in Martinez, M.L. and Psuty, N.P. (eds.) Coastal Dunes: Ecology and Conservation. Ecological Studies, Vol. 171. Berlin: Springer-Verlag, pp. 173–184

Kuroda, R., Kato, M., Tsuge, T. and Aoyama, T. (2021) ‘Arabidopsis phosphatidylinositol 4-phosphate 5-kinase genes PIP5K7, PIP5K8, and PIP5K9 are redundantly involved in root growth adaptation to osmotic stress’, The Plant Journal, 106, pp. 913–927. doi:10.1111/tpj.15207.

Kushawaha, A.K., Khan, A., Sopory, S.K. and Sanan-Mishra, N. (2021) ‘Priming by high temperature stress induces microRNA regulated heat shock modules indicating their involvement in thermopriming response in rice’, Life, 11(4), article 291. doi:10.3390/life11040291.

Laczkó, L., Jordán, S. and Sramkó, G. (2022) ‘The RadOrgMiner pipeline: automated genotyping of organellar loci from RADseq data’, Methods in Ecology and Evolution, 13(9), pp. 1962–1975. doi:10.1111/2041-210X.13937.

Lammers, C., van de Ven, C.N., van der Heide, T. and Reijers, V.C. (2023) ‘Are ecosystem engineering traits fixed or flexible: a study on clonal expansion strategies in co-occurring dune grasses’, Ecosystems, 26(6), pp. 1195–1208. doi:10.1007/s10021-023-00826-4.

Li, H. (2011) ‘A statistical framework for SNP calling, mutation discovery, association mapping and population genetical parameter estimation from sequencing data’, Bioinformatics, 27(21), pp. 2987–2993. doi:10.1093/bioinformatics/btr509.

Li, L., Stoeckert, C.J. Jr and Roos, D.S. (2003) ‘OrthoMCL: identification of ortholog groups for eukaryotic genomes’, Genome Research, 13(9), pp. 2178–2189. doi:10.1101/gr.1224503.

Li, X., Yang, Y., Yao, J., et al. (2009) ‘FLEXIBLE CULM 1 encoding a cinnamyl-alcohol dehydrogenase controls culm mechanical strength in rice’, Plant Molecular Biology, 69, pp. 685–697. doi:10.1007/s11103-008-9448-8.

Lithgow, D., Martínez, M., Gallego-Fernández, J., Hesp, P., Flores, P., Gachuz, S., Rodríguez-Revelo, N., Jiménez-Orocio, O., Mendoza-González, G. and Álvarez-Molina, L. (2013) ‘Linking restoration ecology with coastal dune restoration’, Geomorphology, 199, pp. 214–224. doi:10.1016/j.geomorph.2013.05.007.

Loureiro, J., Rodriguez, E., Doležel, J. and Santos, C. (2007) ‘Two new nuclear isolation buffers for plant DNA flow cytometry: a test with 37 species’, Annals of Botany, 100, pp. 875–888.

Maccaferri, M., Harris, N.S., Twardziok, S.O. et al. (2019) ‘Durum wheat genome highlights past domestication signatures and future improvement targets’, Nature Genetics, 51, pp. 885–895. doi:10.1038/s41588-019-0381-3.

Manel, S., Schwartz, M.K., Luikart, G. and Taberlet, P. (2003) ‘Landscape genetics: combining landscape ecology and population genetics’, Trends in Ecology & Evolution, 18(4), pp. 189–197. 10.1016/S0169-5347(03)00008-9

Maun, M.A. (1998) ‘Adaptations of plants to burial in coastal sand dunes’, Canadian Journal of Botany, 76(5), pp. 713–738. 10.1139/b98-058

Maun, M.A. (2009) The Biology of Coastal Sand Dunes. Oxford: Oxford University Press.

Maun, M.A. and Lapierre, J. (1984) ‘The effects of burial by sand on Ammophila breviligulata’, Journal of Ecology, 72, pp. 827–839. doi:10.2307/2259534.

Meisner, J. and Albrechtsen, A. (2018) ‘Inferring population structure and admixture proportions in low-depth NGS data’, Genetics, 210(2), pp. 719–731. doi:10.1534/genetics.118.301336.

Mendes, F.K., Vanderpool, D., Fulton, B. and Hahn, M.W. (2020) ‘CAFE 5 models variation in evolutionary rates among gene families’, Bioinformatics, 36(2), pp. 551–559. doi:10.1093/bioinformatics/btz782.

Munnik, T. and Vermeer, J.E.M. (2010) ‘Osmotic stress-induced phosphoinositide and inositol phosphate signalling in plants’, Plant, Cell & Environment, 33(4), pp. 655–669. doi:10.1111/j.1365-3040.2009.02097.x.

Nagy, I., Veeckman, E., Liu, C. et al. (2022) ‘Chromosome-scale assembly and annotation of the perennial ryegrass genome’, BMC Genomics, 23, 505. doi:10.1186/s12864-022-08697-0.

Obermayer, R., Leitch, I.J., Hanson, L. and Bennett, M.D. (2002) ‘Nuclear DNA C-values in 30 species double the familial representation in pteridophytes’, Annals of Botany, 90, pp. 209–217.

Pagowski, V. and Micheli, F. (2024) ‘Mind the gap: A review of disjunctions in coastal marine species’, Integrative and Comparative Biology, 64(2), pp. 203–216. 10.1093/icb/icae099

Park, M.Y., Kang, J.-Y. and Kim, S.Y. (2011) ‘Overexpression of AtMYB52 confers ABA hypersensitivity and drought tolerance’, Molecules and Cells, 31(5), pp. 447–454. doi:10.1007/s10059-011-0300-7.

Perrino, E.V., Tomaselli, V., Costa, R. and Pavone, P. (2013) ‘Conservation status of habitats (Directive 92/43 EEC) of coastal and low hill belts in a Mediterranean biodiversity hot spot (Gargano – Italy)’, Plant Biosystems, 147(4), pp. 1006–1028. 10.1080/11263504.2013.860052

Peterson, B.K., Weber, J.N., Kay, E.H., Fisher, H.S. and Hoekstra, H.E. (2012) ‘Double digest RADseq: an inexpensive method for de novo SNP discovery and genotyping in model and non-model species’, PLoS ONE, 7(5), e37135. doi:10.1371/journal.pone.0037135.

Privé, F., Luu, K., Vilhjálmsson, B.J. and Blum, M.G.B. (2020) ‘Performing highly efficient genome scans for local adaptation with pcadapt version 4’, Molecular Biology and Evolution, 37(7), pp. 2153–2154. doi:10.1093/molbev/msaa053.

Ranallo-Benavidez, T.R., Jaron, K.S. and Schatz, M.C. (2020) ‘GenomeScope 2.0 and Smudgeplot for reference-free profiling of polyploid genomes’, Nature Communications, 11, 1432.

Rice, A., Glick, L., Abadi, S., Einhorn, M., Kopelman, N.M., Salman-Minkov, A., Mayzel, J., Chay, O. and Mayrose, I. (2014) ‘The Chromosome Counts Database (CCDB) – a community resource of plant chromosome numbers’, New Phytologist, 206(1), pp. 19–26. doi:10.1111/nph.13191.

Rodríguez-Echeverría, S., Freitas, H. and van der Putten, W.H. (2008) ‘Genetic diversity and differentiation of Ammophila arenaria (L.) Link as revealed by ISSR markers’, Journal of Coastal Research, 24(1), pp. 122–138.

Rozé, F. and Lemauviel-Lavenant, S. (2004) ‘Sand dune restoration in North Brittany, France: a 10-year monitoring study’, Restoration Ecology, 12(1), pp. 29–35. doi:10.1111/j.1061-2971.2004.00264.x.

Samuk, K. (2023) ‘Average nucleotide diversity should be weighted by per-site sample size’, Molecular Ecology Resources, 23(2), pp. 355–358. 10.1111/1755-0998.13738

Skotte, L., Korneliussen, T.S. and Albrechtsen, A. (2013) ‘Estimating individual admixture proportions from next generation sequencing data’, Genetics, 195(3), pp. 693–702. doi:10.1534/genetics.113.154138.

Slatkin, M. (1987) ‘Gene flow and the geographic structure of natural populations’, Science, 236(4803), pp. 787–792. doi:10.1126/science.3576198.

Slaymaker, P., Wresilo, M., Zeltner, D. and Saleh, Y. (2015) ‘Genetic structure of native and restored populations of American beachgrass (Ammophila breviligulata Fern.) along the New Jersey coast’, Journal of Coastal Research, 31(6), pp. 1334–1343. doi:10.2112/JCOASTRES-D-13-00157.1.

Stamatakis, A. (2014) ‘RAxML version 8: a tool for phylogenetic analysis and post-analysis of large phylogenies’, Bioinformatics, 30(9), pp. 1312–1313. doi:10.1093/bioinformatics/btu033.

Stoeckel, S., Grange, J., Fernández-Manjarres, J.F., Bilger, I., Frascaria-Lacoste, N. and Mariette, S. (2006) ‘Heterozygote excess in a self-incompatible and partially clonal forest tree species – Prunus avium L.’, Molecular Ecology, 15(8), pp. 2109–2118. doi:10.1111/j.1365-294X.2006.02926.x.

Sun, J., Lu, F., Luo, Y., Bie, L., Xu, L. and Wang, Y. (2023) ‘OrthoVenn3: an integrated platform for exploring and visualizing orthologous data across genomes’, Nucleic Acids Research, 51(W1), pp. W397–W403. doi:10.1093/nar/gkad313.

Temmerman, S., Meire, P., Bouma, T.J., Herman, P.M.J., Ysebaert, T. and de Vriend, H.J. (2013) ‘Ecosystem-based coastal defence in the face of global change’,

Tkach N, Winterfeld G, Hoffmann MH, Röser M. (2025). Genome size variation in the Poaceae supertribe Poodae, the major grass lineage of temperate climates (tribes Aveneae, Festuceae and Poeae). Plant Systematics and Evolution 311: 24. 10.1007/s00606-025-01949-y

Tutin, T.G. et al. (eds.) (1980) Flora Europaea, Volume 5. Cambridge: Cambridge University Press.

van der Laan, D., van Tongeren, O.F.R., van der Putten, W.H. and Veenbaas, G. (1997) ‘Vegetation development in coastal foredunes in relation to methods of establishing marram grass (Ammophila arenaria)’, Journal of Coastal Conservation, 3, pp. 179–190. 10.1007/BF02905243

Van der Putten, W.H., van Dijk, C. and Troelstra, S.R. (1988) ‘Biotic soil factors affecting the growth and development of Ammophila arenaria’, Oecologia, 76, pp. 313–320.10.1007/BF00379970

Van der Putten, W.H. (1989) Establishment, growth and degeneration of Ammophila arenaria in coastal sand dunes. PhD thesis. Wageningen University and Research

Van der Putten, W.H. (1990) ‘Establishment of Ammophila arenaria (marram grass) from culms, seeds and rhizomes’, Journal of Applied Ecology, 27(1), pp. 188–199. doi:10.2307/2403577.

Van De Walle, R., Massol, F., Vandegehuchte, M.L. and Bonte, D. (2023) ‘The functional composition of dune nematode communities is structured by both biogeographic region and the local marram grass environment’, European Journal of Soil Biology, 117, article 103511. 10.1016/j.ejsobi.2023.103511

Vasimuddin, M., Misra, S., Li, H. and Aluru, S. (2019) ‘Efficient architecture-aware acceleration of BWA-MEM for multicore systems’, in 2019 IEEE International Parallel and Distributed Processing Symposium (IPDPS), pp. 314–324. doi:10.1109/IPDPS.2019.00041.

Wang, T., Tohge, T., Ivakov, A., Mueller-Roeber, B., Fernie, A.R., Mutwil, M., Schippers, J.H. and Persson, S. (2015) ‘Salt-related MYB1 coordinates abscisic acid biosynthesis and signaling during salt stress in Arabidopsis’, Plant Physiology, 169(2), pp. 1027–1041. doi:10.1104/pp.15.00962.

Waterhouse, R.M., Seppey, M., Simão, F.A. et al. (2018) ‘BUSCO applications from quality assessments to gene prediction and phylogenomics’, Molecular Biology and Evolution, 35(3), pp. 543–548. doi:10.1093/molbev/msx319.

Wei, X., Xu, Y., Lyu, L., Xiao, Z., Wang, S., Yang, T. and Jiang, M. (2023) ‘Impacts of ecological restoration on the genetic diversity of plant species: a global meta-analysis’, Journal of Applied Ecology, 60(6), pp. 1149–1160. doi:10.1111/1365-2664.14390.

Whitford, W.G. (2002) Ecology of Desert Systems. San Diego, CA: Academic Press

Williams, S.L. (2001) ‘Reduced genetic diversity in eelgrass transplantations affects both population growth and individual fitness’, Ecological Applications, 11(5), pp. 1472–1488. doi:10.1890/1051-0761(2001)011[1472:RGDIET]2.0.CO;2.

Willis, A.J. and Yemm, E.W. (1961) ‘Braunton Burrows: mineral nutrient status of the dune soils’, Journal of Ecology, 49(2), p. 377. doi:10.2307/2257270.

Zhang, M., Zhang, B., Qian, Q., Yu, Y., Li, R., Zhang, J., Liu, X., Zeng, D., Li, J. and Zhou, Y. (2010) ‘Brittle Culm 12, a dual-targeting kinesin-4 protein, controls cell-cycle progression and wall properties in rice’, The Plant Journal, 63(2), pp. 312–328. doi:10.1111/j.1365-313X.2010.04238.x.

Zhou, C., Brown, M., Blaxter, M., Darwin Tree of Life Project Consortium, McCarthy, S.A. and Durbin, R. (2025) ‘Oatk: a de novo assembly tool for complex plant organelle genomes’, Genome Biology, 26(1), 235. doi:10.1186/s13059-025-03676-6.

