## Supplementary Figures for "Genomic signatures of local adaptation in marram grass highlight the importance of provenance in dune restoration"

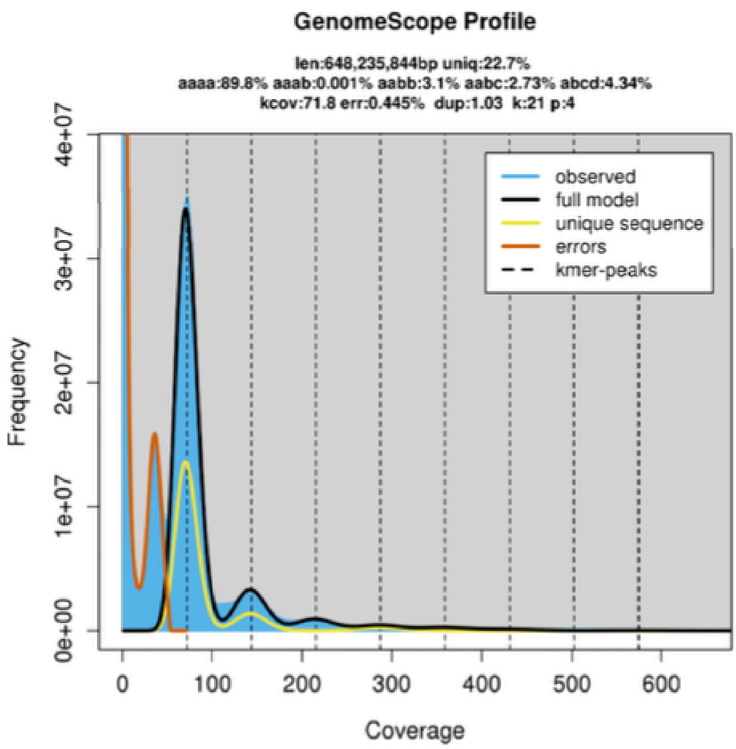


**Figure S1:** K-mer frequency spectrum used to estimate genome size, ploidy, repeat content and heterozygosity. K-mer size = 21 bp; maximum k-mer coverage = 1,000. The distinct low-coverage k-mer peak was interpreted as a genuine genomic signal rather than sequencing errors. This means the overall genome size estimate from GenomeScope is an underestimate. Values above the plot show GenomeScope model estimates, including genome size (*len*), unique sequence (*uniq*), peak k-mer coverage (*kcov*), error rate (*err*), duplication rate (*dup*), k-mer size (*k*) and ploidy (*p*).

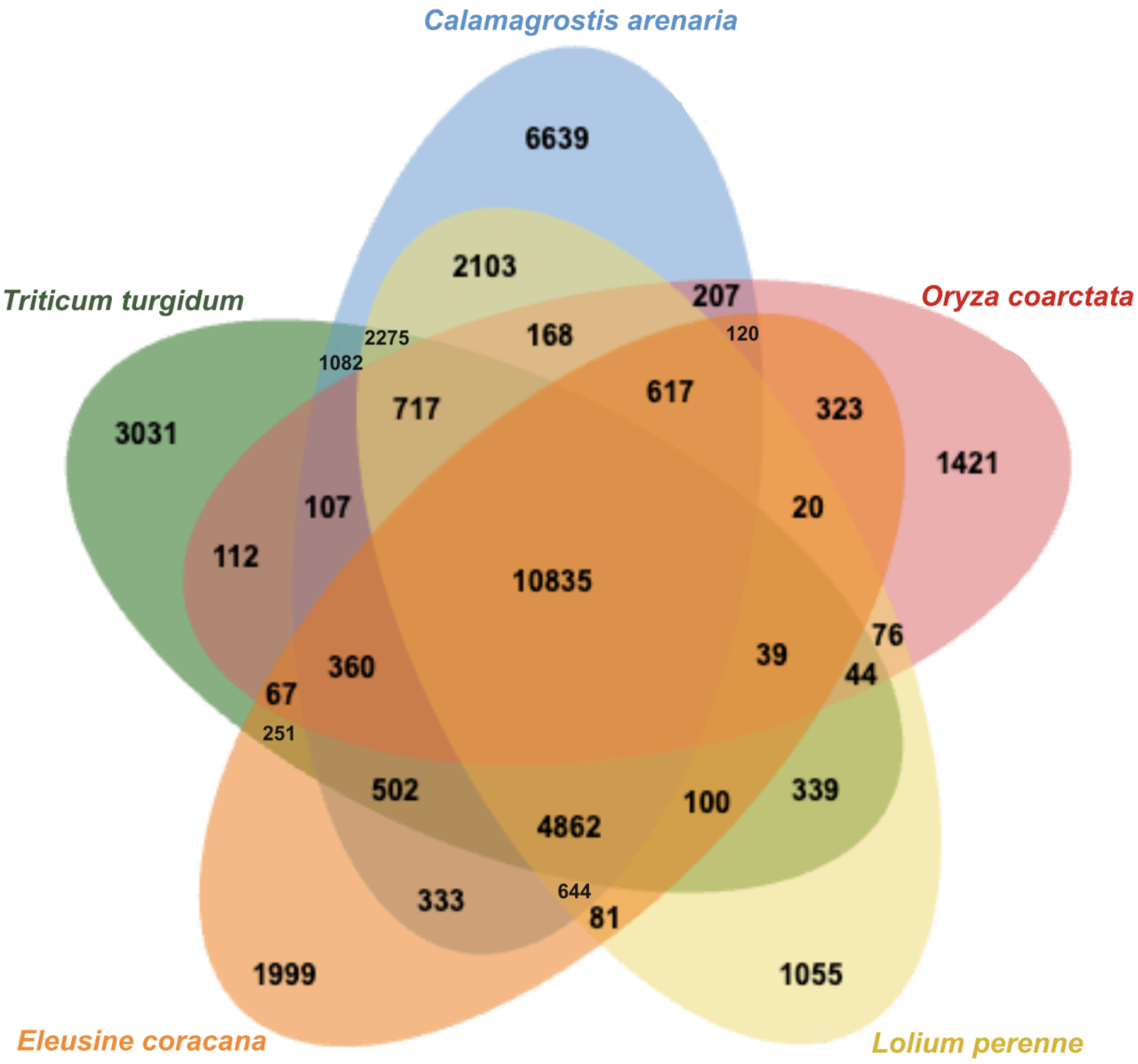


**Figure S2:** Venn diagram showing the number of shared gene orthologous clusters amongst the 4 species included in the analysis on OrthoVenn3.

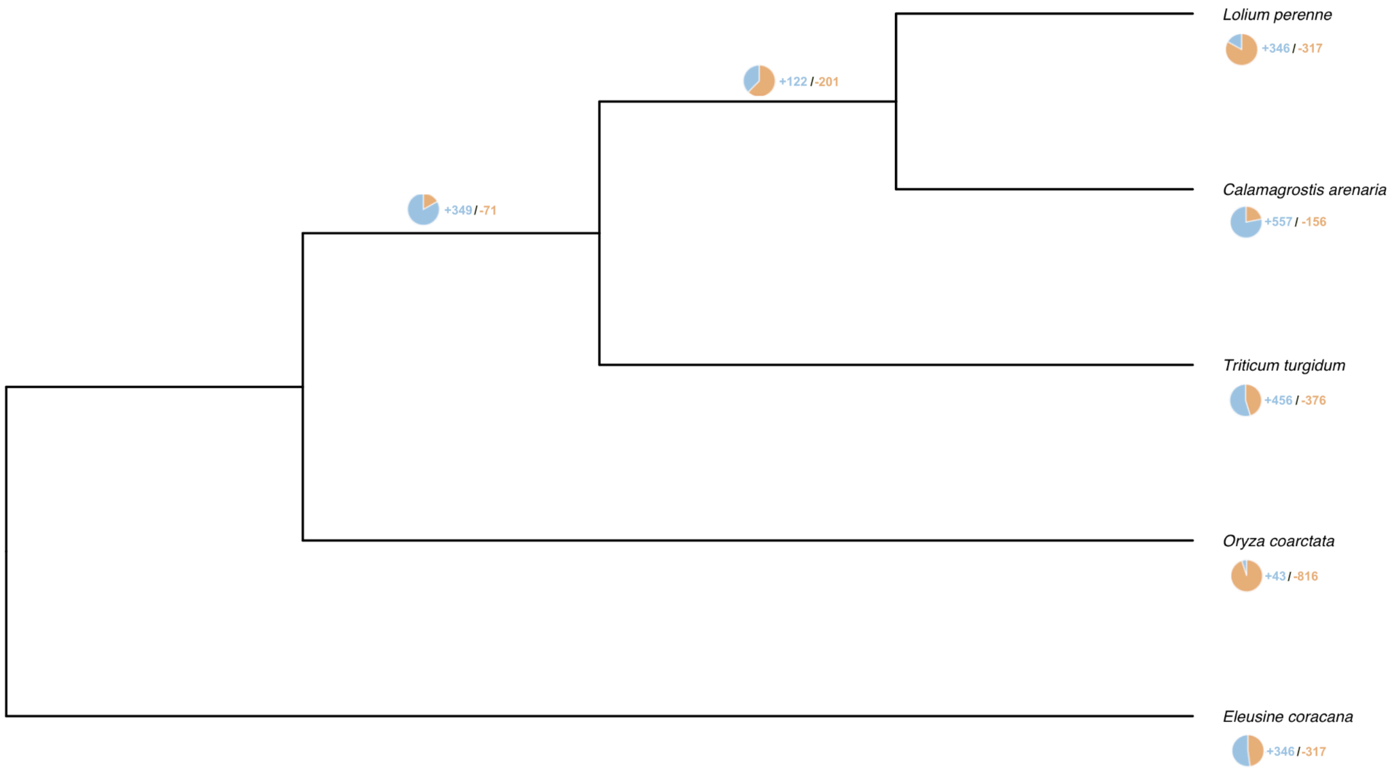


**Figure S3:** Maximum likelihood tree, created using OrthoVenn3, based on single-copy orthologous proteins. The number of gene families that have expanded (blue, +) and contracted (orange, -), as inferred using CAFE5, are represented.

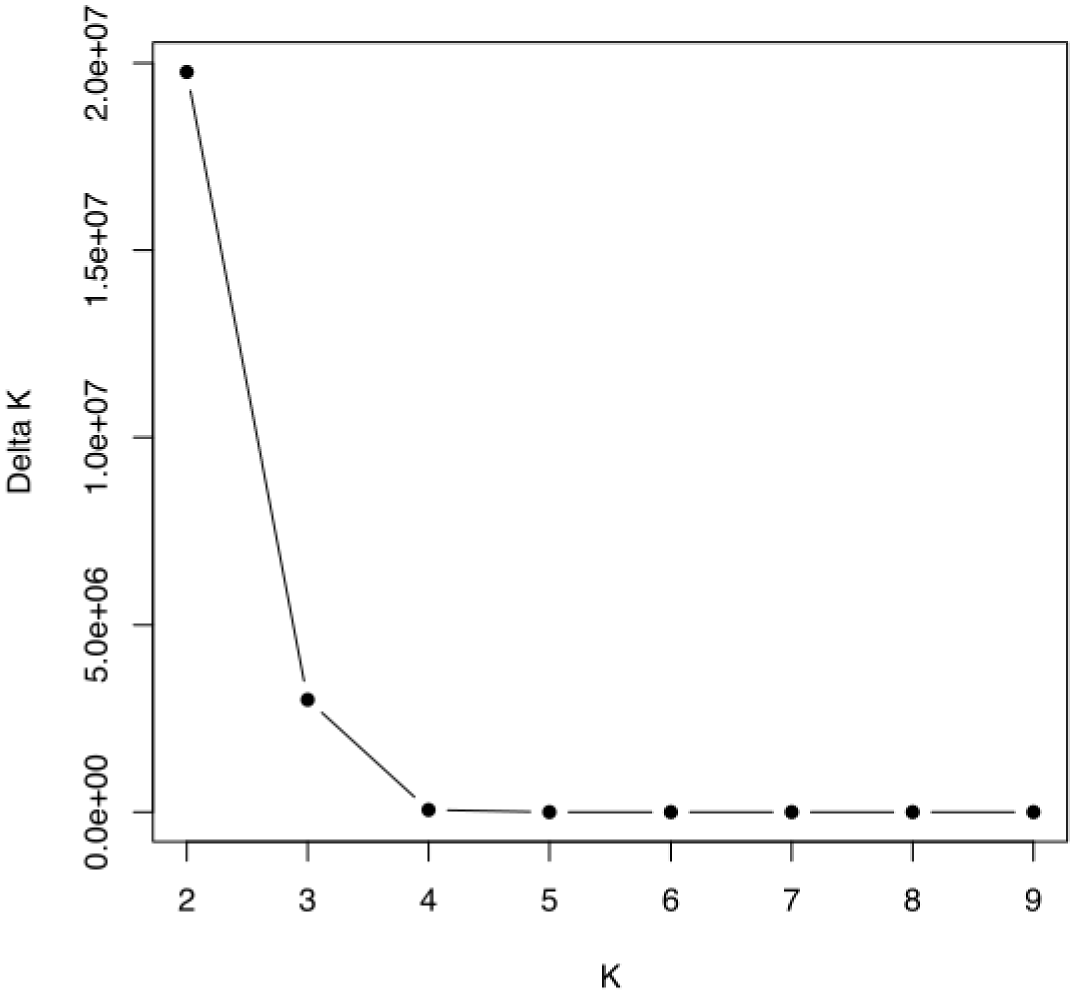


**Figure S4:** Optimal number of genetic clusters in the sampled populations excluding those in Portugal. The delta-K method was used to determine the best value of *K* to describe population structure in the estimated admixture proportions. The highest value of delta-K indicates the optimal value of K.

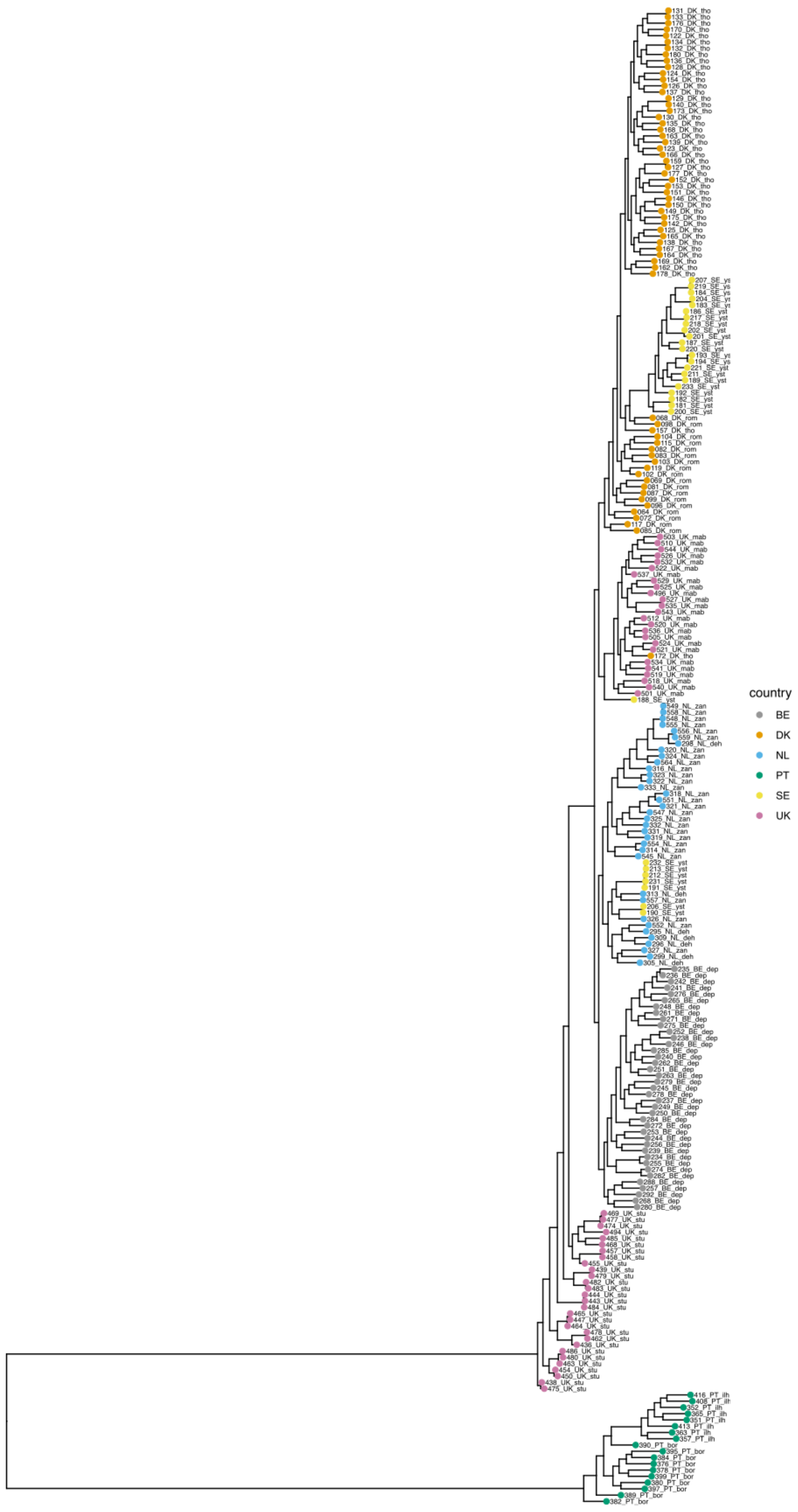


**Figure S5:** Rectangular layout of the phylogenetic tree from Fig. 3A with additional individual level information. Nodes are coloured by country (see key). Numbers reference individual plants and site codes follow the map in Fig. 2A.

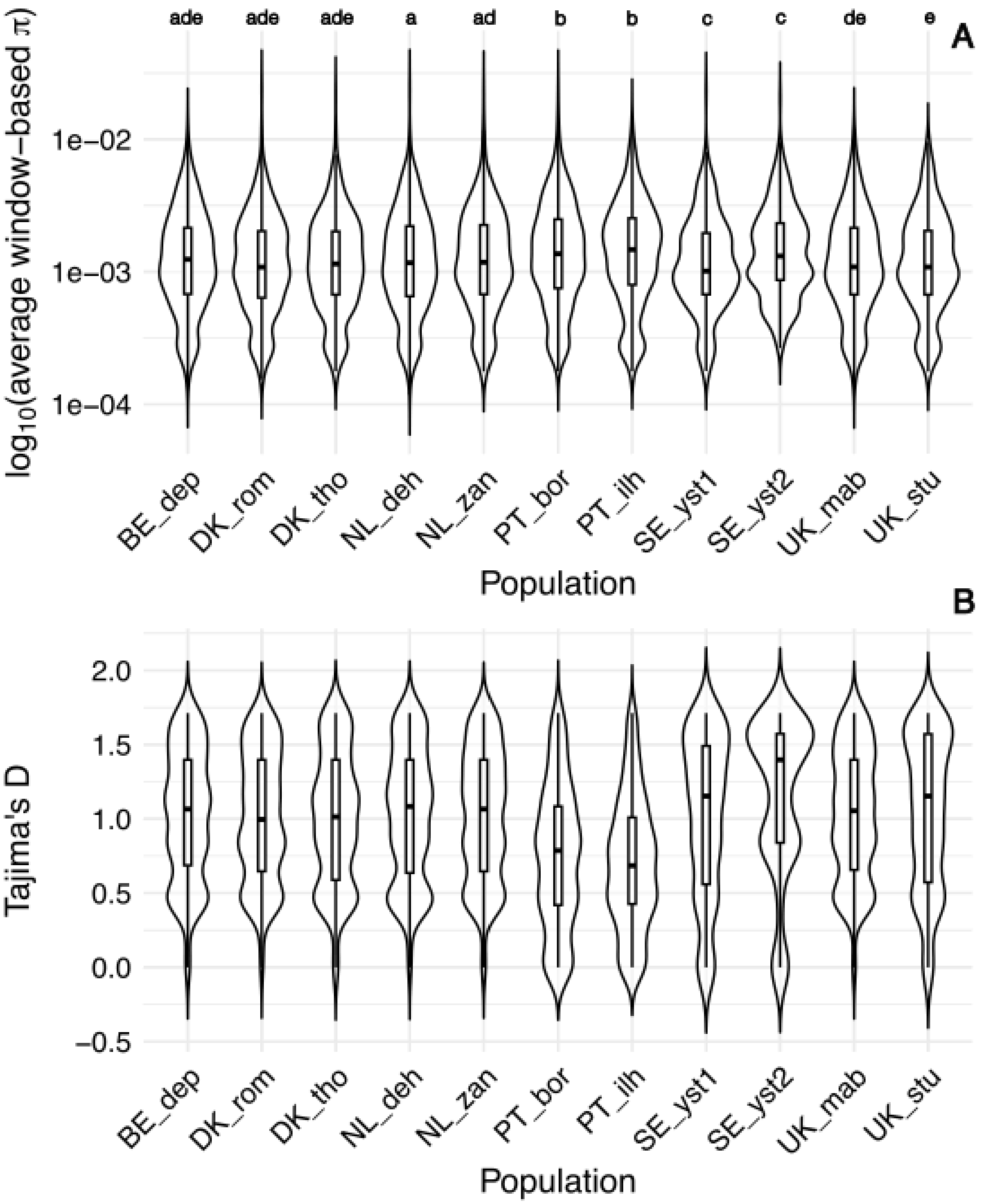


**Figure S6:** Violin plots comparing **A.** nucleotide diversity (**π**). Populations not significantly different (P>= 0.05) following a Dunn's post-hoc test (Table S4) are indicated by sharing the same letter. **B.** Tajima's D across *C. arenaria* populations.

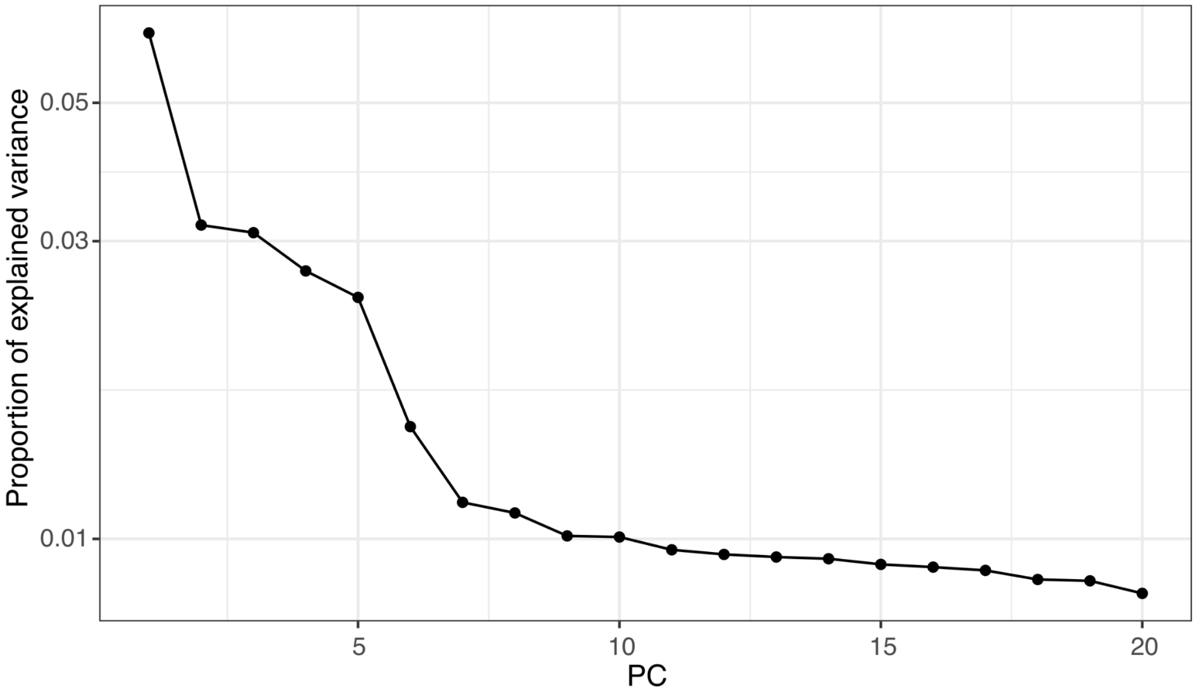


**Figure S7:** Scree plot showing the amount of genetic variation explained by the first 20 principal components, or K clusters. After the first 7 principal components, the proportion of variance explained tails off, so a value of *k* = 7 was selected for the model.
